# SKA2 regulated hyperactive secretory autophagy drives neuroinflammation-induced neurodegeneration

**DOI:** 10.1101/2023.04.03.534570

**Authors:** Jakob Hartmann, Thomas Bajaj, Joy Otten, Claudia Klengel, Anne-Kathrin Gellner, Ellen Junglas, Kathrin Hafner, Elmira A Anderzhanova, Fiona Tang, Galen Missig, Lindsay Rexrode, Katelyn Li, Max L Pöhlmann, Daniel E Heinz, Roy Lardenoije, Nina Dedic, Kenneth M McCullough, Tomasz Próchnicki, Thomas Rhomberg, Silvia Martinelli, Antony Payton, Andrew C. Robinson, Valentin Stein, Eicke Latz, William A Carlezon, Mathias V Schmidt, Chris Murgatroyd, Sabina Berretta, Torsten Klengel, Harry Pantazopoulos, Kerry J Ressler, Nils C Gassen

## Abstract

High levels of proinflammatory cytokines induce neurotoxicity and catalyze inflammation-driven neurodegeneration, but the specific release mechanisms from microglia remain elusive. We demonstrate that secretory autophagy (SA), a non-lytic modality of autophagy for secretion of vesicular cargo, regulates neuroinflammation-mediated neurodegeneration via SKA2 and FKBP5 signaling. SKA2 inhibits SA-dependent IL-1β release by counteracting FKBP5 function. Hippocampal *Ska2* knockdown in mice hyperactivates SA resulting in neuroinflammation, subsequent neurodegeneration and complete hippocampal atrophy within six weeks. The hyperactivation of SA increases IL-1β release, initiating an inflammatory feed-forward vicious cycle including NLRP3-inflammasome activation and Gasdermin D (GSDMD)-mediated neurotoxicity, which ultimately drives neurodegeneration. Results from protein expression and co-immunoprecipitation analyses of postmortem brains demonstrate that SA is hyperactivated in Alzheimer’s disease. Overall, our findings suggest that SKA2-regulated, hyperactive SA facilitates neuroinflammation and is linked to Alzheimer’s disease, providing new mechanistic insight into the biology of neuroinflammation.

## Introduction

Microglia, the resident immune cells of the brain, have critical roles in tissue homeostasis, phagocytic activity and cytokine production. Increasing amounts of pro-inflammatory cytokines, such as IL-1β can be harmful and toxic to neurons and have been associated with neurodegenerative illnesses including Alzheimer’s disease (AD) ^1–4^. However, the specific release mechanisms of pro-inflammatory cytokines from microglia that govern neuroinflammation-driven neurodegeneration are not fully understood.

Secretory autophagy (SA), a non-lytic modality of autophagy for secretion of vesicular cargo involving a stepwise succession of autophagy signaling proteins, cargo receptors and RQ-SNARE fusion proteins, has been linked to peripheral immune responses and inflammation ^5,6^. However, the key molecular mechanisms involved remain elusive and it is unknown whether SA may play a role in neuroinflammation.

Recently, we identified the stress-inducible co-chaperone FK506-binding protein 51 (FKBP5) as a scaffolding protein and key driver of SA that facilitates fusion of the secretory autophagosome with the plasma membrane and subsequent cargo secretion to the extracellular milieu ^7^. FKBP5 promotes the RQ-SNARE complex formation between the secretory autophagosome and the plasma membrane through interaction with several of its key components including vesicle-trafficking protein SEC22B and synaptosomal-associated protein 29 (SNAP29). Interestingly, a scaffolding protein, spindle and kinetochore-associated complex subunit 2 (SKA2), has previously been identified as a potential binding partner of SNAP29 in cervical adenocarcinoma (HeLa S3) cells ^8^, leaving the role of SKA2 in the brain, and its potential involvement in SA, unexplored.

In the current convergent studies, we demonstrate that SA regulates neuroinflammation-mediated neurodegeneration via SKA2 and FKBP5 signaling. SKA2 inhibits SA-dependent IL-1β release by counteracting FKBP5 function in cells, mice and human postmortem brains. Specifically, hyperactivated SA, induced by knockdown of *Ska2* initiates an inflammatory feed-forward vicious cycle resulting in Gasdermin D (GSDMD)-mediated neurotoxicity that ultimately drives neurodegeneration. These results reveal unknown mechanisms and may provide novel targets for intervention and potential prevention of neuroinflammatory and neurodegenerative disorders.

## Results

### SKA2 acts as a molecular roadblock for secretory autophagy that inhibits vesicle-plasma membrane fusion

The final step in SA that allows for cargo secretion (e.g. IL-1β release) is the SNARE complex formation between the R-SNARE SEC22B of the secretory autophagosome and the Q_abc_-SNARE complex, formed by the synaptosomal-associated proteins SNAP23 and SNAP29 and the syntaxins 3 and 4 (STX 3/4) of the plasma membrane. This event leads to the fusion of autophagosome and plasma membrane, with subsequent release of cargo proteins into the extracellular milieu ^6,7,9,10^.

First, we set out to investigate whether SKA2 interacts with components of the SA pathway in the brain. Co-immunoprecipitations (co-IPs) from mouse prefrontal cortex (PFC), hippocampus and amygdala tissue, showed that SKA2 associates with SNAP29 (Fig. 1A), which has previously only been reported in HeLa cells ^8^. In addition, co-IPs revealed associations of SKA2 with other SNARE complex proteins including SEC22B, SNAP23 and STX3 (Fig. 1A). To further examine a potential role for SKA2 in the RQ-SNARE fusion process during SA, we performed co-IPs in a murine microglia cell line (SIM-A9 cells). Knockdown (KD) of *Ska2* enhanced RQ-SNARE complex formation, which was reflected by increased SEC22B binding to SNAP29 as well as SEC22B binding to STX3. Consistent with our previous findings in SH-SY5Y cells ^7^, overexpression (OE) of *Fkbp5* led to a similar increase in binding of SEC22B to SNAP29, as well as SEC22B to STX3 (Fig. 1B-G; unpaired two tailed t-test; SKA2-KD: SEC22B binding to SNAP29, t_6_= 8.945, p < 0.0001, SEC22B binding to STX3, t_6_= 12.94, p < 0.0001; FKBP5-OE: SEC22B binding to SNAP29, t_6_= 6.056, p < 0.001, SEC22B binding to STX3, t_6_= 5.554, p < 0.01; n = 4 per group).

**Figure 1.**
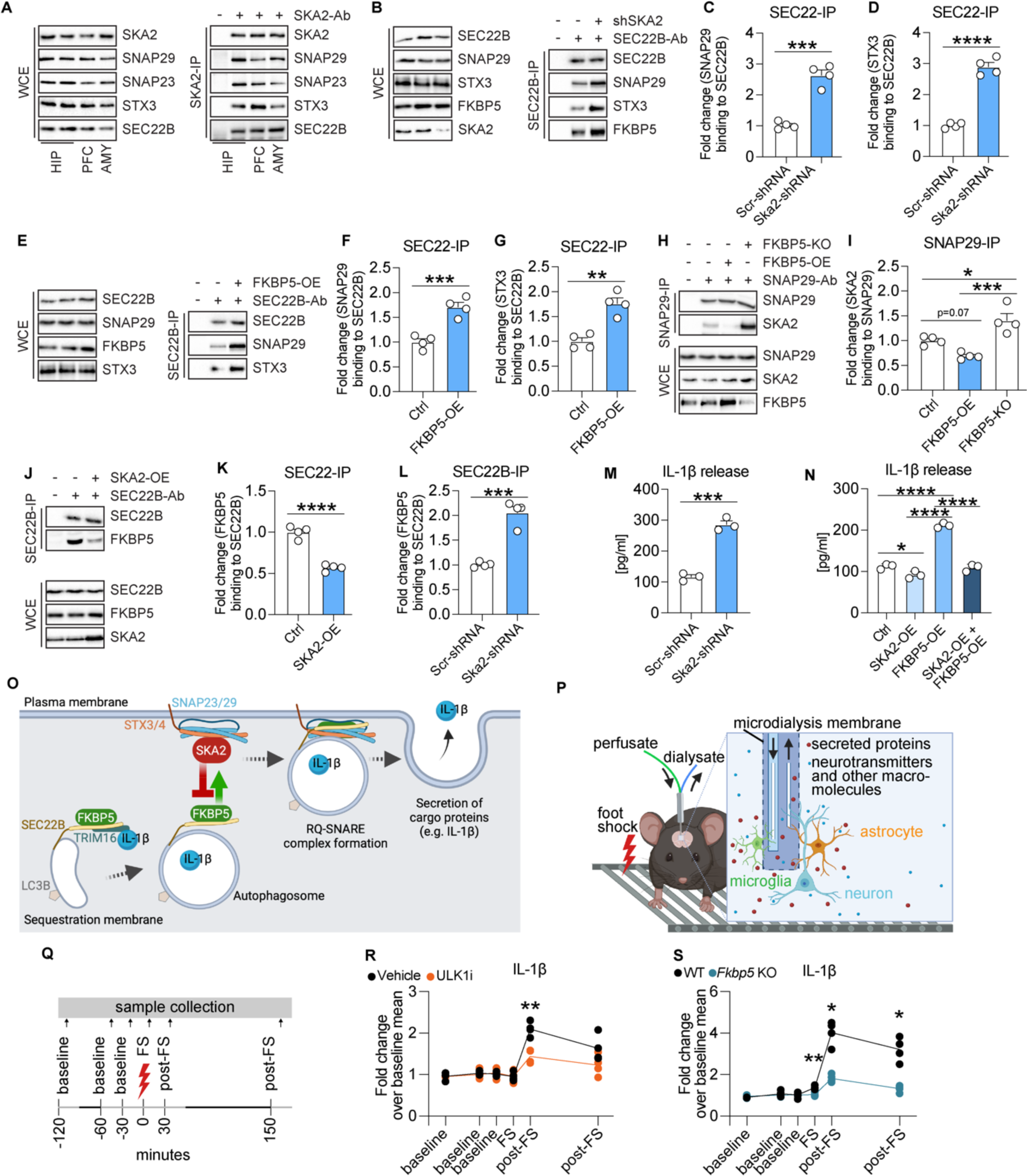
SKA2 and FKBP5 have opposing roles in the final step of secretory autophagy (SA). **(A)** SNAP29, SNAP23, STX3 and SEC22B co-immunoprecipitation (SKA2 IP) and whole cell extract (WCE) in hippocampus (HIP), prefrontal cortex (PFC) and amygdala (AMY) samples of mice (n = 8). **(B-L)** SIM-A9 cells transfected with SKA2, FKBP5 or their respective controls, were harvested 24 h later. After immunoprecipitation (IP) of protein complexes, input and co-IP proteins were quantified by western blotting. **(B, E, H, J)** Representative blots of **(C, D, F, G, I, K and L)**. Graphs display quantification of SNAP29/SEC22B, STX3/SEC22B, SKA2/SNAP29, FKBP5/SEC22B protein interaction after SEC22B or SNAP29 immunoprecipitation (IP). **(M)** and **(N)** IL-1β release from supernatants measured via ELISA 24 h after manipulation of SKA2 and/or FKBP5 expression, and following overnight LPS (100 ng/mL) and treatment with LLOMe (0.25 mM) for 3 h. **(O)** Schematic overview of the SA pathway with SKA2 and FKBP5. The cargo receptor TRIM16, together with SEC22B, transfers molecular cargo (e.g. IL-1β) to the autophagy-related LC3B-positive membrane carriers. SEC22B, now acting as an R-SNARE on the delimiting membrane facing the cytosol, carries out fusion at the plasma membrane in conjunction with the Q_bc_-SNAREs, SNAP23 and SNAP29 (SNAP23/29), and one of the plasma membrane Q_a_-SNAREs, STX3 or STX4 (STX3/4), thus delivering IL-1β to the extracellular milieu, where it exerts its biological functions. FKBP5 acts as a positive regulator of SA by enhancing TRIM16-SEC22B complex formation as well as autophagosome-plasma membrane fusion via the SNARE-protein complex assembly. In contrast, SKA2 inhibits the SNARE-protein complex formation during vesicle-plasma membrane fusion, thereby acting as gatekeeper of SA. **(P-Q)** Schematic overview of *in vivo* microdialysis and the experimental design and timeline; each sample was collected over 30 min indicated by the light grey lines. Quantifications of IL-1β, determined by capillary-based immunoblotting from *in vivo* medioprefrontal cortex microdialysis of C57Bl/6NCrl mice injected intraperitoneally with ULK1 inhibitor (ULK1i, an autophagy inhibitor) or saline **(R)** as well as of wild type (WT) and global *Fkbp5* knockout mice **(S) (**n = 4 mice per group). FS foot shock. Unpaired, two tailed t-test for simple comparisons, one-way analysis of variance (ANOVA) + Tukey’s post hoc test, repeated measures two-way ANOVA + Šidák’s multiple comparisons post hoc test; * = p < 0.05; ** = p < 0.01; *** = p < 0.001; **** = p < 0.0001. Data are presented as mean + SEM. n = mean derived from 3-4 independent *in vitro* experiments.

Next, we tested a functional link between SKA2 and FKBP5, and found that knockout (KO) of *Fkbp5* led to a significant increase in SKA2 to SNAP29 binding, while *Fkbp5* overexpression had the opposite effect (Fig. 1H and I; one-way ANOVA, F_2, 9_ = 17.28, p < 0.001; Tukey’s post hoc test: ctrl vs. FKBP5-OE, p = 0.07, ctrl vs. FKBP5-KO, p < 0.05, FKBP5-OE vs. FKBP5-KO, p < 0.001). Importantly, co-IPs revealed that overexpression of *Ska2* significantly reduced FKBP5 to SEC22B binding (Fig. 1J and K; unpaired two tailed t-test; t_6_= 10.27, p < 0.0001), while a *Ska2* KD induced the opposite effect (Fig. 1B and L; unpaired two tailed t-test; t_6_= 8.140, p < 0.001). Together, these data demonstrate that SKA2 is directly involved in RQ-SNARE complex formation and appears to regulate SA in opposition to the role of FKBP5.

IL-1β is a well-established cargo protein released via SA ^6^. In order to investigate whether SKA2 alters IL-1β secretion, we manipulated *Ska2* expression in (Lipopolysaccharide (LPS) and L-leucyl-L-leucine methyl ester (LLOMe)-primed) SIM-A9 cells and analyzed the supernatants with enzyme-linked immunosorbent assay (ELISA). *Ska2* KD significantly increased release of IL-1β, 24 h after transfection (Fig. 1M; unpaired two tailed t-test; t_4_= 11.99, p < 0.001). In addition, overexpression of *Ska2* resulted in reduced IL-1β release, while *Fkbp5* overexpression led to the opposite effect. Strikingly, *Ska2* overexpression was able to reverse the increase in IL-1β release induced by *Fkbp5* overexpression (Fig. 1N; one-way ANOVA, F_3, 8_ = 158.6, p < 0.0001; Tukey’s post hoc test: ctrl vs. SKA2-OE, p < 0.05, ctrl vs. FKBP5-OE, p < 0.0001, SKA2-OE vs. FKBP5-OE, p < 0.0001, FKBP5-OE vs. SKA2 + FKBP5 OE, p < 0.0001). Together these results suggest that SKA2 and FKBP5 play contrasting roles in the final step of SA, in particular with regards to IL-1β release. While FKBP5 enhances the formation of the RQ-SNARE complex and subsequent IL-1β release, SKA2 decreases it, thereby acting as gatekeeper of this secretory pathway (Fig. 1O).

## Activation of SA increases IL-1β release *in vivo*

To confirm that secretion of IL-1β is dependent on the autophagic machinery *in vivo*, we assessed its extracellular dynamics in medial PFC (mPFC) using *in vivo* microdialysis in C57Bl/6NCrl mice injected with the selective ULK1 inhibitor (ULK1i) MRT68921, an established blocker of early autophagy machinery. Microdialysates were collected under baseline conditions and following acute and strong, foot-shock stress with the aim to potentiate IL-1β release. We previously showed that acute stress increases activity of SA and subsequent cargo release ^7^ (Fig. 1P-Q). There were no changes in IL-1β secretion under baseline conditions between the treatment groups. In contrast, stress-induced IL-1β release was significantly decreased in mice treated with ULK1i compared to vehicle controls (Fig. 1R; repeated measures two-way ANOVA, time x treatment interaction: F_5, 30_ = 7.064, p < 0.001; Šidák’s multiple comparisons post hoc test, p < 0.01; n = 4 per group). Along these lines, acute stress induced a significant increase in IL-1β secretion in wild type mice, an effect that was blunted in *Fkbp5* KO mice (Fig. 1S; repeated measures two-way ANOVA, time x genotype interaction: F_5, 30_ = 34.15, p < 0.0001; Šidák’s multiple comparisons post hoc test, p < 0.05; n = 4 per group). These data further validate our *in vitro* findings of IL-1β secretory regulation. They also underline the importance of the SA pathway on brain physiology and the potential impact on neuroinflammation.

## Hyperactivity of SA leads to NLRP3 inflammasome activation, neuroinflammation-induced neurodegeneration

In order to better understand the relevance of SA and its impact on brain physiology, we performed a viral-mediated shRNA-dependent KD of *Ska2* in the hippocampus of C57Bl/6J mice. Remarkably, KD of *Ska2* (Fig. S1A) induced pronounced neurodegeneration compared to viral infection with a scrambled control shRNA. *Ska2* KD resulted in complete hippocampal atrophy within six weeks of the viral injection (Fig. 2A). This was also reflected in decreased expression of the neuronal marker NeuN and drastically reduced CA1 thickness, two- and four-weeks following KD of *Ska2* (Fig. 2B; paired t-test: 2 weeks, t_4_= 3.194, p < 0.05; 4 weeks, t_3_= 6.711, p < 0.01). In addition, immunohistochemistry (IHC) with IBA1 revealed an increase in microglia numbers in the hippocampus, 2 weeks after viral-mediated KD of *Ska2*, an effect that was even more pronounced after 4 weeks (Fig. 2C; paired t-test: 2 weeks, t_4_ = 4.295, p < 0.05; 4 weeks, t_3_ = 7.165, p < 0.01). Moreover, expression of the astrocyte marker GFAP was increased 2 and 4 weeks following *Ska2* KD in the hippocampus (Fig. 2D; paired t-test: 2 weeks, t_4_= 5.524, p < 0.01; 4 weeks, t_3_= 5.764, p < 0.05). No assessment of marker expression was possible at week 6 due to complete hippocampal atrophy. Importantly, the neurodegenerative process and inflammatory response were not caused by off-target effects since a similar phenotype was observed upon KD of *Ska2* with a second *Ska2*-shRNA, targeting a different region of the gene and packaged into a different viral capsid serotype (Fig. S1B and C). Taken together, these findings indicate that hippocampal disruption of SKA2 leads to progressive neuroinflammation and subsequent neurodegeneration, likely through an overactivation of the SA pathway.

**Figure 2.**
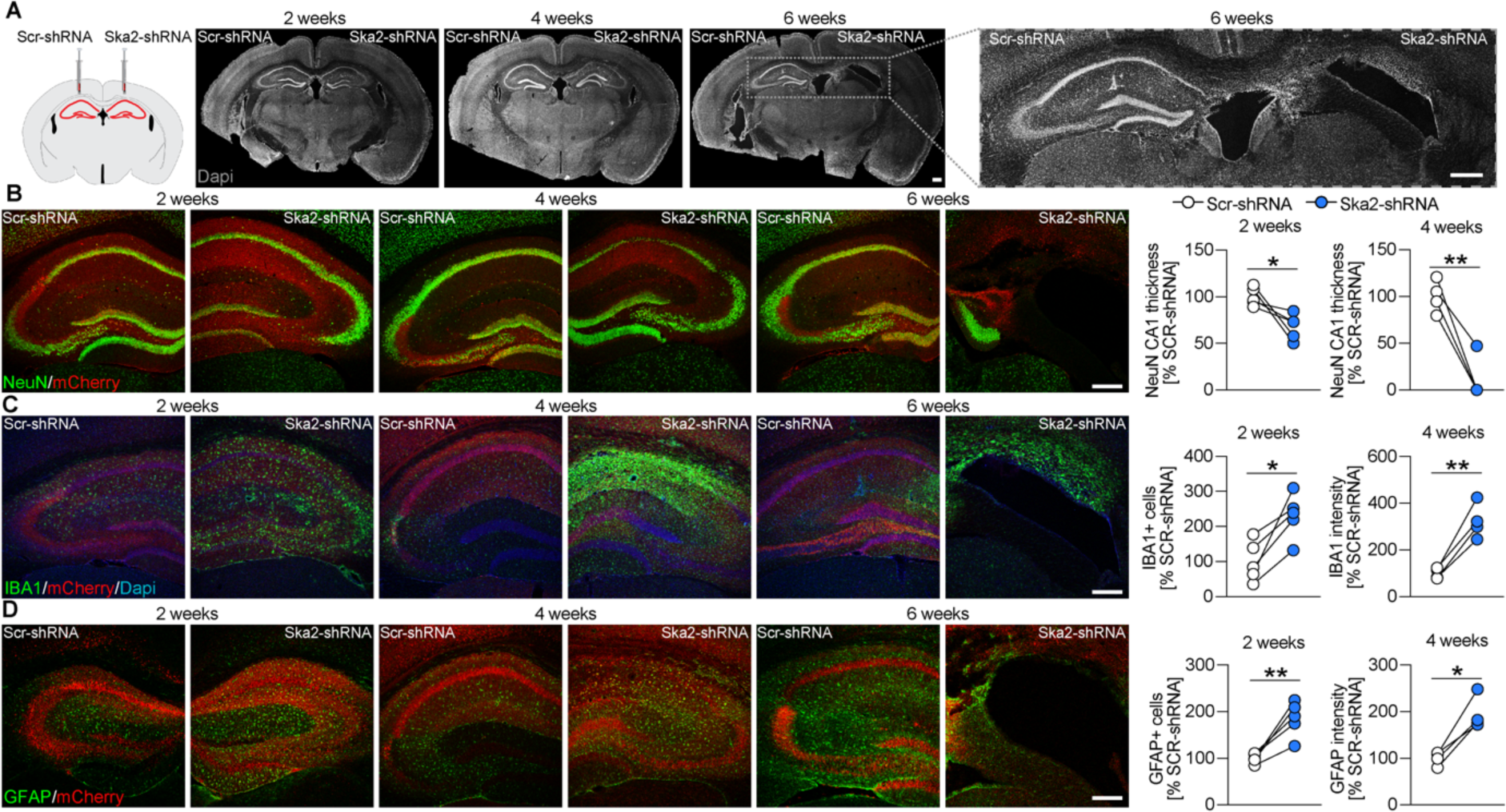
Hippocampal *Ska2* knockdown induces neuroinflammation-mediated neurodegeneration in mice. **(A)** Schematic representation of viral injections (Scr-shRNA-AAV and Ska2-shRNA-1-AAV) into the hippocampus (left). (right) Representative IHC images of DAPI (gray) 2, 4 and 6 weeks after viral injections. **(B)** IHC images of NeuN (green) and mCherry (red, viral marker) 2, 4 and 6 weeks after viral injection. Quantification of CA1 thickness 2 and 4 weeks after viral injection. **(C)** IHC images of IBA1 (green), mCherry (red) and DAPI (blue) 2, 4 and 6 weeks after viral injection. Quantification of IBA1 expression 2 and 4 weeks after viral injection. **(D)** IHC images of GFAP (green) and mCherry (red) 2, 4 and 6 weeks after viral injection. Quantification of GFAP expression 2 and 4 weeks after viral injection. Paired t-test: * = p < 0.05; ** = p < 0.01; n = 4 to 5 mice per time point. Scale bars represent 250 µm.

Previously, increasing intensities of pro-inflammatory stimuli (e.g. microbial components or endogenous cytokines) have been shown to induce sequential activation of vesicular and Gasdermin D (GSDMD)-mediated IL-1β secretory pathways ^11^. In order to investigate whether altered SA activity (and thus IL-1β release) is able to modulate inflammasome formation, we used a clonal inflammasome reporter overexpressing fluorescently tagged ASC (apoptosis-associated speck-like protein containing a CARD) ^12^ in wild type and *Sec22b* KO SIM-A9 cells. Already under control conditions, inhibition of SA activity (through *Sec22b* KO) resulted in significantly less ASC specks compared to wild type controls (Fig. 3A; unpaired two tailed t-test: t_4_ = 3.206, p < 0.05). Moreover, ASC specks were significantly increased in SIM-A9 WT cells following LPS treatment or KD of *Ska2* compared to vehicle or Scr-shRNA controls (Fig. 3B; 2-way ANOVA: main LPS treatment effect, F_1,31_ = 10.60, p < 0.01, main *Ska2* knockdown effect, F_1,31_ = 5.482, p < 0.05). In contrast, the LPS- and *Ska2* KD-dependent inflammasome formation was abolished when the SA pathway was disrupted in *Sec22b* KO SIM-A9 cells Fig. 3C; 2-way ANOVA: n. s.). In order to identify which inflammasome is stimulated through increased activity of SA, we investigated protein lysates of organotypic hippocampal slice cultures. KD of *Ska2* resulted in significantly increased binding of SEC22B to SNAP29, reflective of enhanced SA activity (Fig. 3D; unpaired two tailed t-test: t_4_ = 4.113, p < 0.01). The kinase NEK7 is an important requirement in the activation of the NLRP3 (NOD-, LRR- and pyrin domain-containing protein 3) inflammasome via NLRP3-NEK7 association^13^. Along these lines, NEK7 binding to NLRP3 was significantly increased following *Ska2* KD (Fig. 3E, unpaired two tailed t-test: t_4_ = 2.998, p < 0.05). Therefore, we next investigated whether KD of *Ska2* in the hippocampus of mice and thus overactivation of SA, may serve as an inflammasome-inducing signal leading to GSDMD-mediated IL-1β secretion. Indeed, KD of *Ska2* led to increased ASC expression and ASC specks formation (Fig. 3F-G; paired t-test: 2 weeks, ASC+ cells, t_2_ = 6.414, p < 0.05, ASC specks, t_2_ = 6.937, p < 0.05; 4 weeks, ASC+ cells, t_2_ = 8.511, p < 0.05, ASC specks, t_2_ = 10.99, p < 0.01) as well as CASPASE-1 (CASP-1) expression (Fig. 3H-I; paired t-test: 2 weeks, t_3_ = 2.842, p = 0.06; 4 weeks, t_3_ = 3.367, p < 0.05), indicative of inflammasome activation. Inflammasome-activated CASP-1 cleaves GSDMD to release the N-terminal domain which forms pores on the membrane that enable passage of cytokines including IL-1β ^14,15^. Accordingly, the expression levels of full length (FL) GSDMD as well as its cleaved N-terminal domain (GSDMD N-term) were increased at 2 weeks following KD of *Ska2* (Fig. 3J-K; unpaired two tailed t-test; GSDMD FL/ β-actin: t_18_= 4.105, p < 0.001; GSDMD N-term/GSDMD FL: t_18_= 9.259, p < 0.0001). Together, these data provide significant mechanistic evidence that hyperactivated SA (through KD of *Ska2*) is able to create an inflammatory feed-forward vicious cycle resulting in a GSDMD-mediated excessively neurotoxic environment to ultimately catalyze neuroinflammation and neurodegeneration (Fig. 3L).

**Figure 3.**
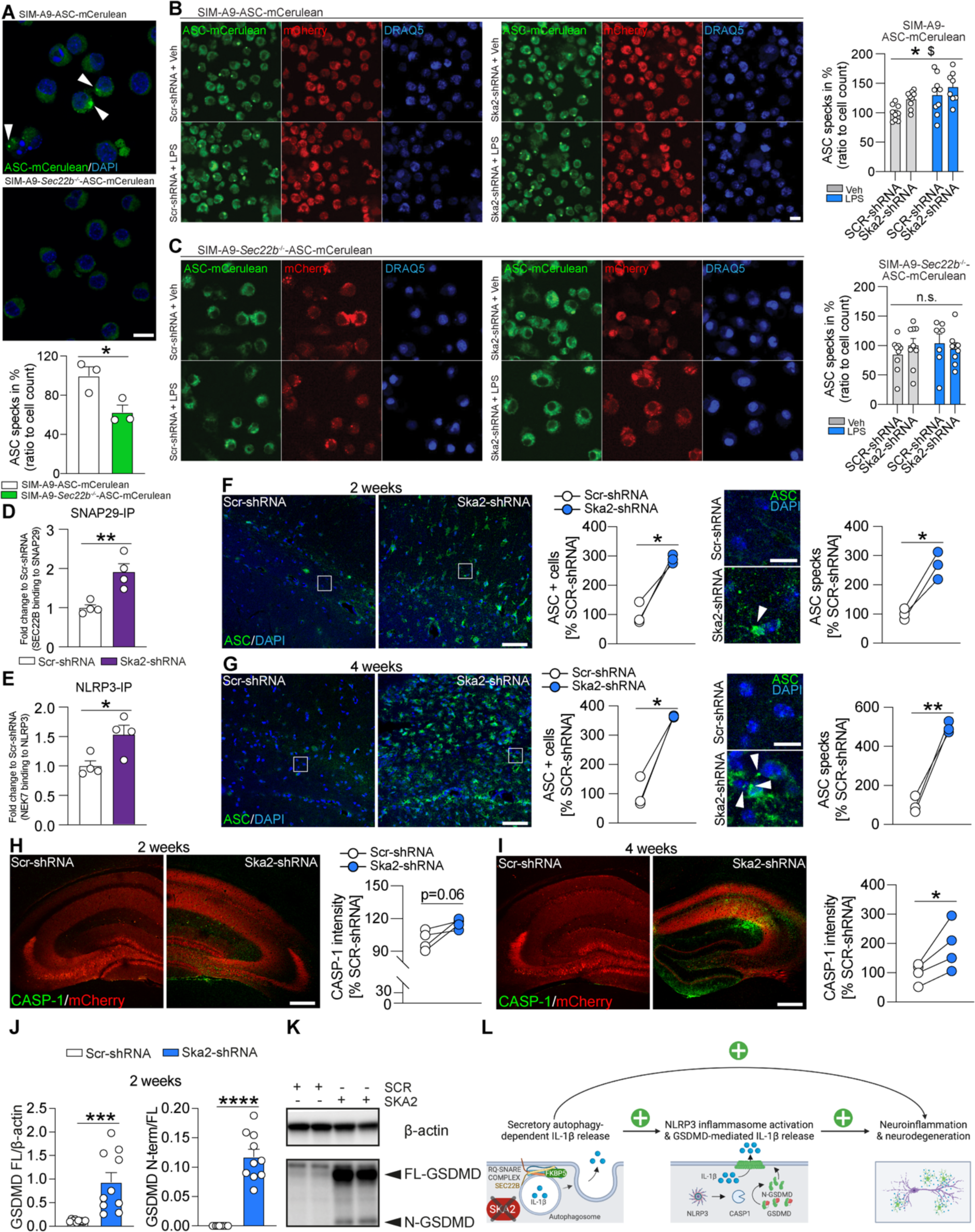
Hyperactivity of SA induces inflammasome formation. **(A)** SIM-A9 *Sec22b^-/-^* cells expressing ASC (apoptosis-associated speck-like protein containing a CARD) - mCerulean (via epifluorescence) show a significantly decreased number of intracellular (white arrows) ASC-specks compared to wild type (WT) SIM-A9 cells (n = mean derived from 3 independent *in vitro* experiments). **(B)** In WT SIM-A9 cells (n = mean derived from 2 independent *in vitro* experiments, with n = 7 to 9 replicates corresponding to wells of a 96-well plate) knockdown of *Ska2* or LPS treatment leads to a significantly increased number of intracellular ASC-specks compared to Scr-shRNA or LPS-treated cells (two-way analysis of variance (ANOVA): main Ska2 effect (* = p < 0.05), main LPS effect ($ = p < 0.05)). **(C)** In contrast, knockdown of *Ska2* or LPS treatment does not have any effects on the number of ASC-specks in SIM-A9 *Sec22b^-/-^* cells (n = mean derived from 2 independent *in vitro* experiments, with n = 7 to 9 replicates corresponding to wells of a 96-well plate). **(D-E)** Knockdown of *Ska2* leads to significantly increased SEC22B binding to SNAP29 as well as NEK7 binding to NLRP3 in protein lysates of organotypic hippocampal slice cultures (n = 4 per group). (**F**) IHC images of ASC (green) and DAPI (blue) 2 weeks after viral injection (Scr-shRNA-AAV and Ska2-shRNA-1-AAV) into the hippocampus. Quantification of ASC+ cells (left) and ASC specks (right) 2 weeks after viral injection. **(G)** IHC images of ASC (green) and DAPI (blue) 4 weeks after viral injection (Scr-shRNA-AAV and Ska2-shRNA-1-AAV) into the hippocampus. Quantification of ASC+ cells (left) and ASC specks (right) 4 weeks after viral injection. **(H)** IHC images of CASPASE-1 (CASP-1) (green) and mCherry (red, viral marker) 2 weeks after viral injection (Scr-shRNA-AAV and Ska2-shRNA-1-AAV) into the hippocampus (left). (right) Quantification of CASP-1 expression 2 weeks after viral injection. **(I)** IHC images of CASP-1 (green) and mCherry (red, viral marker) 4 weeks after viral injection (Scr-shRNA-AAV and Ska2-shRNA-1-AAV) into the hippocampus (left). (right) Quantification of CASP-1 expression 4 weeks after viral injection. **(J)** Full length Gasdermin D (GSDMD FL) levels as well as the ratio of the cleaved N-terminal form of GSDMD (GSDMD N-term) to GSDMD FL are increased 2 weeks after *Ska2* knockdown. **(K)** Examples blots of (E). **(L)** Schematic overview of the interaction between secretory autophagy (SA) and the GSDMD-mediated IL-1β release. SKA2 depletion results in increased SA-dependent IL-1β release, serving as a molecular vicious feed-forward loop for inflammasome activation. Inflammasome assembly activates CASP-1 enzymatic function. ASC in the inflammasome complex recruits CASP-1. Activation of CASP-1 cleaves GSDMD to release the N-terminal domain, which forms pores in the plasma membrane for uncontrolled IL-1β release. Paired or unpaired, two tailed t-test for simple comparisons, two-way analysis of variance: * = p < 0.05; ** = p < 0.01; *** = p < 0.001, **** = p < 0.0001. Data are presented as mean + SEM. Scale bar represents 5 µm in A, 50 µm in F-G (left), 10 µm in B, F-G (right) and 250 µm in H-I.

## *Ska2* knockdown in the hippocampus leads to cognitive impairment

The severe hippocampal atrophy observed at 4 weeks following *Ska2* KD resulted in expected spatial memory (Y-maze) and novel object recognition memory impairments in mice (Fig. 4A-B; Y-maze, 2-way ANOVA: condition x arm interaction, F_2,48_ = 3.626, p < 0.05, Tukey’s post hoc test: familiar arm A vs. novel arm, p < 0.001, familiar arm B vs. novel arm, p < 0.01; n = 9 per group; novel object test, unpaired t-test: t_15_ = 2.840, p < 0.05; n = 9 Scr-shRNA group, n = 8 Ska2-shRNA group). The observed cognitive deficits were not accompanied by changes in general locomotor activity (open field test; unpaired t-test: p > 0.05, n = per group) or anxiety-related behavior (elevated plus maze (EPM); unpaired t-test: p > 0.05, n = 8 Scr-shRNA group, n = 9 Ska2-shRNA group), which confound learning and memory tasks (Fig. 4C-D). These findings indicate that hippocampal disruption of SKA2 leads to cognitive impairment.

**Figure 4.**
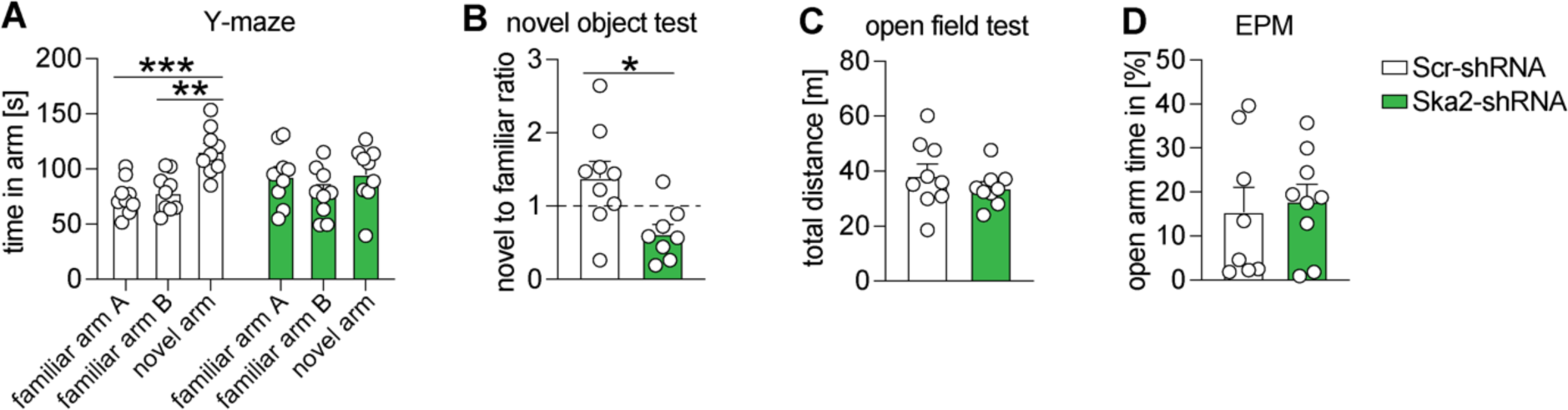
Hippocampal *Ska2* knockdown leads to cognitive impairment in mice. **(A)** In the Y-maze test, mice injected with Scr-shRNA spent significantly more time in the novel arm compared to both familiar arms (A and B). These effects were abolished following *Ska2* knockdown (n = 9 per group). **(B)** In contrast to control animals, Ska2-shRNA mice did not discriminate between a novel and familiar object during the novel object recognition test (n = 9 Scr-shRNA group, n = 8 Ska2-shRNA group). **(C-D)** *Ska2* knockdown did not alter general locomotor activity (p>0.05) in the open field test or anxiety-related behavior (p>0.05) in the elevated plus maze (EPM), n = 8-9 per group. Unpaired, two tailed t-test for simple comparisons, two-way analysis of variance + Tukey’s post hoc test: * = p < 0.05; ** = p < 0.01; *** = p < 0.001. Data are presented as mean + SEM.

## Secretory autophagy is increased in human postmortem Alzheimer’s disease samples

Given the impact that SA and its regulators, SKA2 and FKBP5, have on brain function in mice, we continued to explore the relevance of this secretory pathway and its components in the human brain. To investigate the relationship of the *SKA2* and *FKBP5* genes with phenotypic traits, we searched these loci in the Atlas of genome-wide association studies (GWAS) Summary Statistics (http://atlas.ctglab.nl/PheWAS) ^16^. Interestingly, Phenome-Wide Association Studies (PheWAS) associated the *FKBP5* locus with, among others, immunological traits such as lymphocyte count, white blood cell count and monocyte percentage of white cells. PheWAS of the *SKA2* locus associated with cognitive as well as with immunological traits, including intelligence and cognitive performance as well as monocyte percentage of white cells, granulocyte percentage of myeloid white cells and monocyte count (Fig. S2A-B, Table S1-S2).

Next, performing co-IPs, we confirmed an association of SKA2 with SNAP29 in human PFC, amygdala and hippocampus in postmortem tissue from healthy subjects (Fig. 5A, Table S3). IHC of brain sections from healthy human subjects (n = 5) revealed a pronounced expression of SKA2 in the adult hippocampus (mid body coronal sections consisting of the dentate gyrus and the stratum oriens and pyramidal cell layers of the CA1, CA2, CA3 and CA4 subregions). Additional morphological and co-expression analyses revealed a prominent expression of SKA2 not only in pyramidal neurons, but also in microglia (Fig. 5B-D, Table S4).

**Figure 5.**
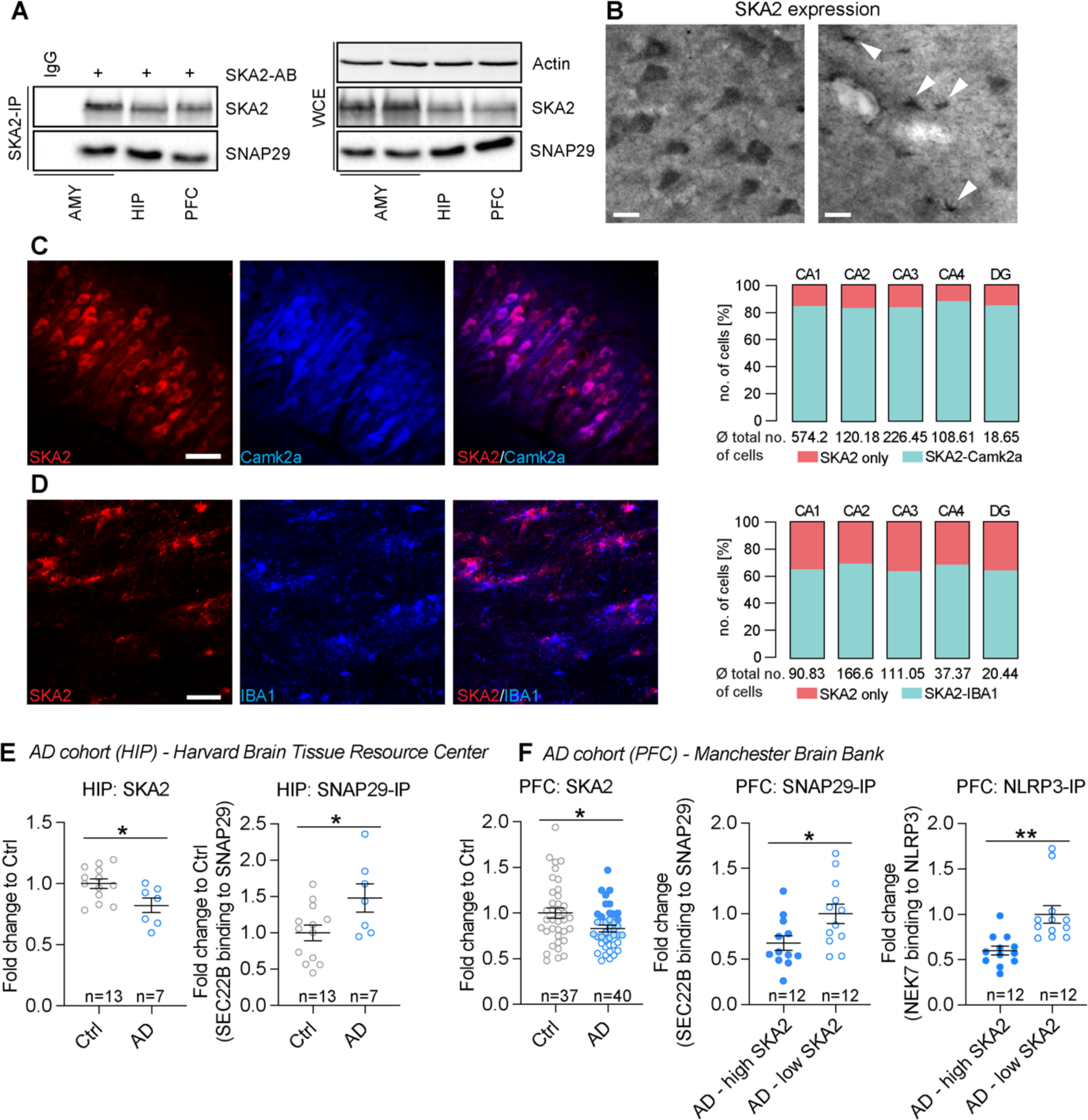
Secretory autophagy is increased in human postmortem Alzheimer’s disease samples. **(A)** SNAP29 co-immunoprecipitation (SKA2 IP) and whole cell extract (WCE) control in amygdala (AMY), hippocampus (HIP) and prefrontal cortex (PFC) human postmortem samples (n = 3). **(B)** SKA2 immunostaining in neurons in CA1 stratum pyramidale (left) and microglia in CA1 stratum oriens (right) of the HIP from control subjects. **(C)** Representative co-immunohistochemistry (IHC) image (left) and quantification (right) of SKA2 (red) and neuronal marker Camk2a (blue) in the HIP (n = 5). **(D)** Representative co-IHC image in stratum oriens CA1 of the HIP (left) and quantification (right) of SKA2 (red) and microglia marker IBA1 (blue) (n = 5). **(E)** AD cohort from the Harvard Brain Tissue Resource Center (n = 13 (Ctrl), 7 (AD)): SKA2 protein expression (left) is significantly decreased in the hippocampus of AD subjects while SEC22B binding to SNAP29 (right) is significantly increased in hippocampus tissue of AD subjects. **(F)** AD cohort from the Manchester Brainbank (n = 37 (Ctrl), 40 (AD)): SKA2 protein expression (left) is significantly decreased in the prefrontal cortex (PFC) of AD subjects. SEC22B binding to SNAP29 (middle) as well as NEK7 binding to NLRP3 (right) is significantly increased in PFC tissue of the top 12 low compared to the top 12 high SKA2 expressing AD subjects. ANCOVA: * = p < 0.05; data are presented as mean + SEM. Scale bars represent 50 µm for D and F, 100 µm for E.

Given that our data suggest a critical role for SKA2 in SA and neuroinflammation-induced neurodegeneration, we further investigated whether a hyperactivated SA pathway is involved in the pathophysiology of AD. Therefore, we analyzed SKA2 protein expression using Western blotting, and performed co-IPs and subsequent capillary-based immune analysis to explore SEC22B to SNAP29 binding in the hippocampus of a cohort of AD cases (n = 7) and age matched controls (n = 13) (Table S5). SKA2 expression was significantly decreased in AD (Fig. 5E, left; ANCOVA: F_1,19_ = 6.9123, p < 0.05), while SEC22B to SNAP29 binding was increased (Fig. 5E, right; ANCOVA: F_1,19_ = 5.6769, p < 0.05). Importantly, we were able to validate these findings in an independent replication cohort of prefrontal cortex samples of AD cases (n = 40) and age matched controls (n = 37) (Table S6), demonstrating significantly reduced SKA2 expression in AD (Fig. 5F, left; ANCOVA: F_1,76_ = 6.4994, p < 0.05), and thus pointing towards hyperactivated SA in AD. Along these lines, SEC22B binding to SNAP29 was significantly increased in AD cases with the lowest SKA2 expression (n = 12) compared AD cases with the highest SKA2 expression (n = 12) (Fig. 5F, middle; ANCOVA: F_1,23_ = 2.411, p < 0.05). Moreover, NEK7 binding to NLRP3 was significantly increased in the low SKA2 expression compared to the high SKA2 expression AD group (Fig. 5F, right; ANCOVA: F_1,23_ = 3.696, p < 0.01), indicative of augmented NLRP3 inflammasome activation.

Collectively, our data suggest an important role of SA and its regulators, FKBP5 and SKA2, in microglia and brain function. Importantly, we provide evidence for an involvement of SA in inflammasome activation, neuroinflammation and the pathophysiology of AD.

## Discussion

There is a growing body of evidence that secretory autophagy (SA) may be implicated in processes ranging from cancer to cell death and degeneration, due to its diverse cargo ranging from granule content to cytokines ^17–21^. Moreover, a decrease in lysosomal integrity, which is a hallmark of SA ^6,7,22^, might subsequently reduce the function of homeostatic neuroprotective lytic autophagy. Along these lines, our results support a model in which SA differentially regulates neuroinflammation-induced neurodegeneration via SKA2 and FKBP5 signaling and is implicated in AD.

Overactivation of this pathway in mice through viral-mediated KD of hippocampal *Ska2* resulted in strong microglial activation and recruitment, leading to complete hippocampal atrophy within 6 weeks of viral injection. IL-1β is an essential cytokine, but its release needs to be strictly controlled to avoid severe inflammatory manifestations. Several pathways have been proposed to mediate its release involving secretory lysosomes, exosomes, micro-vesicles and autophagic vesicles as well as GSDMD-dependent routes ^5,11,23^. Further, it has been suggested that pathways that involve the translocation of IL-1β into intracellular vesicles of lysosomal origin (that eventually fuse with the plasma membrane) are primarily in control of IL-1β release upon low pro-inflammatory stimuli, whereas stronger stimulation or concomitant cell stress induces uncontrolled secretion of IL-1β via the GSDMD-mediated pathway ^11,22^.

Our data suggest that hyperactivated SA, in its most severe form, represents a strong enough stimulus to result in a vicious molecular feed-forward loop that triggers the production and uncontrolled secretion of pro-inflammatory cytokines through GSDMD-mediated pathways, ultimately leading to pyroptosis and neurodegeneration. Interestingly, *SKA2* DNA methylation has been linked to a decrement in thickness of the PFC ^24^ and less SKA2 expression in surrounding tissue ^25^.

Clinically, AD is characterized by several features, notably a progressive cognitive decline involving loss of memory and higher executive functioning ^26^. Excessive SA in mice, which was induced via KD of *Ska2,* resulted not only in severe hippocampal neuroinflammation and neurodegeneration, but also in cognitive impairment. Intriguingly, PheWAS identified immunological and cognitive traits such as monocyte count, intelligence and cognitive performance with the *SKA2* locus as well as immunological phenotypes such as lymphocyte count with the *FKBP5* locus. Moreover, the *FKBP5* variant rs1360780 has been associated with altered cognitive function in aged individuals ^27^.

Importantly, our human postmortem data suggest that markers of SA activity are increased in the hippocampus and prefrontal cortex of AD brains (i.e. decreased SKA2 expression along with enhanced SEC22B to SNAP29 binding). Along these lines, increased FKBP5 expression has previously been linked to AD in several brain regions, and higher FKBP5 levels were associated with AD progression ^28^. Human genetic studies have identified microglia as a key cell type governing the risk for AD ^1,2^. Notably, *FKBP5* mRNA expression is increased in microglia of entorhinal cortex postmortem samples from individuals with AD ^29^. Together, these results provide further evidence for the involvement of SA and its key regulators, SKA2 and FKBP5, in cognitive function and AD pathology.

There is increasing evidence from epidemiological and preclinical studies for the effects of environmental factors including early-life and chronic stress as well as traumatic experiences on microglia biology, which in turn might affect an individual’s susceptibility to neurodegenerative diseases ^2,30–32^. However, the underlying molecular mechanisms that mediate the crosstalk between neuronal stress circuits and the immune system remain largely unclear. Previous studies suggest that stress-exposure may precipitate disease risk by increasing inflammation in the periphery and in the brain ^33–36^. Mechanistically, the effects of stress on neuroinflammation, and ultimately disease risk, could be mediated by stress-responsive genes and pathways able to modulate immune function. Indeed, the stress-inducible protein FKBP5 has been to shown to contribute to NF-κB-driven inflammation ^37^. Notably, we have recently demonstrated that dexamethasone and glucocorticoid-mediated stress enhance SA via FKBP5, thereby driving extracellular BDNF maturation and synaptic plasticity as well as elevated immune signaling ^7^. In the current study, our data reveal that SA-dependent and stress-induced release of IL-1β is impaired in the mPFC of *Fkbp5* KO mice. This puts SA in a prime position to mediate the crosstalk between neuronal stress circuits and the immune system. Thus, in the brain, depending on SA’s activity level and specific cargo, this pathway might be involved in the entire spectrum of processes ranging from synaptic plasticity during learning and memory to neuroinflammation-induced neurodegeneration in the pathophysiology of diseases such as AD.

SKA2 expression was also shown to be regulated by glucocorticoids and involved in glucocorticoid receptor (GR) signaling. However, in contrast to FKBP5, SKA2 expression is decreased following dexamethasone treatment and SKA2 is suggested to enhance GR translocation to the nucleus in A549 human lung epithelial cells ^38^. Thus, chronic or severe traumatic stress might lead to increased FKBP5 expression and decreased SKA2 levels, thereby increasing the activity of SA, which in the long run may precipitate in neurotoxicity and neurodegeneration.

Multiple lines of evidence support the pathogenic role of neuroinflammation in psychiatric illness. Elevated levels of central and peripheral cytokines have been detected in individuals with childhood trauma and stress-related psychiatric disorders ^39–41^. Notably, single nucleotide polymorphism and epigenetic marks within the *FKBP5* and *SKA2* genes have repeatedly been associated with stress-related psychiatric diseases including major depressive disorder (MDD) and PTSD as well as suicide risk ^24,25,42–45^. This is interesting considering that psychiatric illnesses such as MDD and PTSD can increase the risk for dementia and AD ^46–48^.

In summary, this study identifies SKA2 as a novel and crucial molecular roadblock of SA in the mammalian brain. Our work highlights the central role of SA in the regulation of inflammasome activation and neuroinflammation-induced neurodegeneration, as well as its implication in the pathophysiology of AD.

## Methods

## Neuro-2a cells

N2a cells (ATCC, CCL-131) were maintained under standard conditions in Dulbecco’s Modified Eagle Medium (DMEM) supplemented with 10% FBS and 1% antibiotic-antimycotic (all Thermo Fisher Scientific) at 37°C and 5% CO_2_ (vol/vol). For cell culture experiments, cells were seeded in 24 well plates at 35,000 cells/well. Transfection was performed the next day using Lipofectamine 2000 Transfection Reagent (Thermo Fisher Scientific), following the manufacturer’s protocol. Cells were harvested 48 h post transfection using TrypLE Express (Thermo Fisher Scientific).

## SIM-A9 cells

The murine microglia cell lines SIM-A9 wild type (Kerafast, END001), SIM-A9 *Sec22b* KO and SIM-A9 *Fkbp5* KO ^7^ were cultured at 37°C, 6% CO_2_ in DMEM high glucose with GlutaMAX (Thermo Fisher Scientific, 10566016), supplemented with 10% FBS (Thermo Fisher, 10270-106), 5% horse serum (Thermo Fisher Scientific, 16050-122) and 1% antibiotic-antimycotic (Thermo Fisher Scientific, 15240-062). With 1x trypsin-EDTA (Thermo Fisher Scientific, 15400-054) detached SIM-A9 cells (2 × 10^6^) were resuspended in 100 μl of transfection buffer [50 mM HEPES (pH 7.3), 90 mM NaCl, 5 mM KCl, and 0.15 mM CaCl_2_]. Up to 2 μg of plasmid DNA was added to the cell suspension, and electroporation was carried out using the Amaxa 2b-Nucleofector system (Lonza). Cells were replated at a density of 10^50^ cells/cm^2^.

### Animals & animal housing

Male mice, aged 2 to 4 months, were used for all experiments. For experiments in wild type animals, C57BL/6J mice were obtained from The Jackson Laboratory (Bar Harbor, ME, USA). For *in vivo* brain microdialysis experiments, C57BL/6NCrl mice (Martinsried, Germany) as well as global *Fkbp5^-/-^* ^49^ mice and respective wild type controls (Martinsried, Germany) were used. All animals were kept under standard laboratory conditions and were maintained on a 12 h light–dark cycle (lights on from 0700 to 1900 h), with food and water provided *ad libitum*. All experiments conformed to National Institutes of Health guidelines and were carried out in accordance with the European Communities’ Council Directive 2010/63/EU and the McLean Hospital Institutional Animal Care and Use Committee. All efforts were made to minimize animal suffering during the experiments. The protocols were approved by the committee for the Care and Use of Laboratory animals of the Government of Upper Bavaria, Germany or by the local Institutional Animal Care and Use Committee, respectively.

### Preparation of organotypic hippocampal slice cultures (OHSC)

Neonatal Thy1-GFP M mice with a sparse expression of green fluorescent protein (GFP) in principal neurons in cortex and hippocampus ^50^, Jackson Laboratory Stock #007788) were used. Pups aged between P7-9 (sex not determined) were decapitated, brains removed, and hippocampi isolated from both hemispheres in ice-cold 1x minimum essential medium (MEM) with EBSS, 25 mM HEPES and 10 mM Tris buffer (pH 7.2) supplemented with Penicillin (100 I.U./ml) and Streptomycin (100 µg/ml). Hippocampi were cut into coronal slices (thickness 350 µm) using a tissue chopper (McIlwain). Slices were transferred onto “confettis” Millipore biopore membrane, ∼3×5 mm) which were placed on semiporous Millicell-CM inserts (0.4 µm pore size; Merck-Millipore). The inserts were put into cell culture dishes (35mm, 4 slices/dish). OHSC were cultured according to the interface method ^51^ in 1 ml medium per dish at pH 7.2, 35 °C and a humidified atmosphere with 5% CO_2_. Culture medium contained 0.5 × MEM with EBSS and 25 mM HEPES, 1mM L-Glutamine, 25% Hanks’ Balanced Salt solution (HBSS), 25% heat-inactivated horse serum, Penicillin (100 I.U./ml) and Streptomycin (100 µg/ml; all Fisher Scientific). The medium was changed one day after preparation and every other day afterwards. Knockdown experiments were performed between 13-15 days in culture (DIC). Medium was harvested and snap frozen in liquid nitrogen directly. OHSCs were lysed in T PER^TM^ Tissue Extraction Reagent (Thermo Fisher, 78510), supplemented with protease (Sigma, P2714) and phosphatase (Roche, 04906837001) inhibitor cocktails.

### Human studies

Tissue blocks were obtained from the Harvard Brain Tissue Resource Center / NIH NeuroBioBank (HBTRC/NBB), McLean Hospital, Belmont, MA, USA. *Healthy control subjects:* Tissue blocks containing the hippocampus, prefrontal cortex (Brodmann area 9) and the amygdala from donors with no history of neurologic or psychiatric conditions were used for histochemical and immunocytochemical investigations as well as for immunoprecipitation analyses. See Table S9-S10 for details of all subjects. All brains underwent a neuropathological examination including several brain regions. These studies did not include subjects with evidence for gross and/or macroscopic brain changes, or clinical history, consistent with cerebrovascular accident or other neurological disorders. Subjects with Braak stages III or higher (modified Bielchowsky stain) were not included. None of the subjects had significant history of substance dependence within 10 or more years from death, as further corroborated by negative toxicology reports. Absence of recent substance abuse is typical for samples from the HBTRC, which receives exclusively community-based tissue donations. Postmortem diagnoses were determined by two clinicians on the basis of retrospective review of medical records and extensive questionnaires concerning social and medical history provided by family members. *Alzheimer’s disease discovery cohort:* Tissue blocks containing the hippocampus from donors with Alzheimer’s disease (n = 7) and healthy control subjects (n = 13) were used for western blotting and co-immunoprecipitation. All subjects were characterized clinically and neuropathologically as above. ‘Control’ cases had Braak & Braak scores of 0-II, sparse plaque pathology, and were rated as having low probability of AD. AD cases had Braak & Braak scores of III-VI and were rated as having intermediate or high probability of AD. Neither group presented with additional relevant neuropathological findings. Groups were matched based on demographic factors (Table S5).

#### Alzheimer’s disease replication cohort

Fresh, frozen tissue was taken from the superior frontal gyrus (Brodmann area 8) of the frontal cortex from 77 brains (AD: n = 40, Ctrl: n = 37) of donors who were participants of a large prospective cognitive aging cohort known as the University of Manchester Longitudinal Study of Cognition in Normal Healthy Old Age Cohort (UMLCHA)^52,53^. Samples were used for western blotting. Samples were acquired through the Manchester Brain Bank with ethical approval granted from the Manchester Brain Bank Committee. AD neuropathology was determined as described above. Groups were matched based on demographic factors (Table S6).

### Plasmids

#### shRNA Construction

shRNA plasmids against mSka2 were constructed as follows: A shRNA plasmid containing a U6 promoter and a multiple cloning site followed by a mCherry gene driven by the PGK promoter was purchased from VectorBuilder Inc (Santa Clara, CA). Target sequences for mSka2 were derived from https://www.sigmaaldrich.com/life-science/functional-genomics-and-rnai/shrna/individual-genes.html. We designed custom 58nt oligos with AgeI/EcoRI restriction sites, annealed them to generate double stranded DNA fragments and ligated this fragment into the AgeI/EcoRI sites of pshRNA to generate Ska2-shRNA-1 (Ska2-shRNA-1: 5’ CGAGAGGATCGTGATGCATTTCTCGAGAAATGCATCACGATCCTCTCG 3’) and Ska2-shRNA-2 (Ska2-shRNA-2: 5’ ACTGATACCCAGCATTCATTTCTCGAGAAATGAATGCTGGGTATCAGT 3’). Similar, a scrambled control was constructed (Scrambled-shRNA sequence: 5’ CCTAAGGTTAAGTCGCCCTCGCTCGAGCGAGGGCGACTTAACCTTAGG 3’). Restriction digest and Sanger Sequencing confirmed the resulting plasmids.

#### mSka2 overexpression

The plasmid overexpressing Ska2 (EF1A>mSka2[NM_025377.3]:IRES:EGFP) and its control (EF1A>EGFP) were purchased from VectorBuilder Inc (Santa Clara, CA).

#### Fkbp5 overexpression

pRK5-FKBP5-FLAG have been described previously ^54^.

### Immunoblotting analysis

Frozen human brain tissue was pulverized on dry ice using a pre-cooled mortar and pestle, then transferred to an ice-cold homogenizer on ice. Mice were sacrificed by decapitation in the morning (08:00 to 08:30 am) following quick anesthesia by isoflurane. Brains were removed, snap-frozen in isopentane at −40°C, and stored at −80°C until further processing. Tissue punches of the prefrontal cortex, hippocampus and amygdala were collected. Protein extracts from cell lines, mouse brains or pulverized human postmortem brains were obtained by lysing in T PER^TM^ Tissue Extraction Reagent (Thermo Fisher, 78510) or lysis radio-immuno precipitation (RIPA) buffer, supplemented with protease (Sigma, P2714) and phosphatase (Roche, 04906837001) inhibitor cocktails. Samples were sonicated and heated at 95 °C for 10 min if necessary. Proteins were separated by SDS–polyacrylamide gel electrophoresis (PAGE) and electro-transferred onto nitrocellulose membranes. Blots were placed in Tris-buffered saline solution supplemented with 0.05% Tween (Sigma, P2287; TBS-T) and 5% non-fat milk for 1 h at room temperature and then incubated with primary antibody (diluted in TBS-T) overnight at 4 °C. Subsequently, blots were washed and probed with the respective horseradish-peroxidase- or fluorophore-conjugated secondary antibody for 1 h at room temperature. The immuno-reactive bands were visualized either using ECL detection reagent (Millipore, WBKL0500) or directly by excitation of the respective fluorophore. Recording of the band intensities was performed with the ChemiDoc MP and corresponding ImageLab software from Bio-Rad or the Odyssey CLx system interfaced with Image Studio version 4.0. Microdialysates obtained from *in vivo* acute stress experiments were analyzed using capillary-based immunoassays (Jess, ProteinSimple) and IL1B (1:50, Gene Tex, GTX74034) antibody.

#### Quantification

All protein data were normalized to ACTIN, GAPDH or VCP, which was detected on the same blot.

#### Primary antibodies used

FKBP5 (1:1000, Bethyl, A301-430A), ACTIN (1:5000, Santa Cruz Biotechnology, sc-1616), GAPDH (1:8000, Millipore CB1001), SNAP29 (1:1000, Sigma, SAB1408650), SNAP23 (1:1000, Sigma, SAB2102251), STX3 (1:1000, Sigma, SAB2701366), SEC22B (1:1000, Abcam, ab181076), SKA2 (1:1000, Thermo Fisher, PA5-20818), SKA2 (1:500, Millipore-Sigma, SAB3500102) GSDMD (1:1000, Cell Signaling Technology, 39754), VCP (1:10000, Abcam, Ab11433), NEK7 (1:50, Abcam, Ab133514), NLRP3 (1:50, Cell Signaling Technology, 15101).

#### Secondary antibodies used

anti-rabbit-IgG (1:1000, Cell Signaling, 7074), anti-mouse-IgG (1:1000, Cell Signaling, 7076), IRDyes 800CW donkey anti-Rabbit (1:20,000, LI-COR Biosciences, 926-32213), IRDye 680RD goat-anti-mouse (1:20,000, LI-COR Biosciences, 926-68070).

### Co-immunoprecipitation

For immunoprecipitation, cells were cultured for 3 days after transfection. Cells were lysed in Co-IP buffer [20 mM tris-HCl (pH 8.0), 100 mM NaCl, 1 mM EDTA, and 0.5% Igepal complemented with protease (Sigma) and phosphatase (Roche, 04906837001) inhibitor cocktail] for 20 min at 4 °C with constant mixing. The lysates (from cells or brain tissue as described above) were cleared by centrifugation, and the protein concentration was determined and adjusted (1.2 mg/ml); 1 ml of lysate was incubated with 2.5 μg of SEC22B, SNAP29 or SKA2 antibody overnight at 4°C with constant mild rotating. Subsequently, 20 μl of bovine serum albumin (BSA)-blocked protein G Dynabeads (Invitrogen, 100-03D) were added to the lysate-antibody mix followed by a 3 h incubation at 4 °C. Beads were washed three times with phosphate buffered saline (PBS), and bound proteins were eluted by adding 60 μl of Laemmli sample buffer and by incubation at 95°C for 5 min. 5 to 15 μg of the input lysates or 2.5 to 5 μl of the immunoprecipitates were separated by SDS–PAGE and analyzed by western blotting. Immunoprecipitates of protein extracts obtained from human post mortem brains were analyzed using capillary-based immunoassays (Jess, ProteinSimple). When quantifying co-immunoprecipitated proteins, their signals were normalized to input protein and to the precipitated interactor protein.

### ELISA

The solid-phase sandwich ELISA (enzyme-linked immunosorbent assay) for the following antibody detection was performed according to the manufacturer’s protocol: IL-1β (Thermo Fisher, BMS6002). Briefly, microwells were coated with mouse antibody followed by a first incubation with biotin-coupled anti mouse antibody, a second incubation with streptavidin-HRP and a final incubation with the SIM-A9 culture medium. Amounts of respective proteins were detected with a plate reader (iMARK, Bio-Rad) at 450 nm.

### RNA extraction and qPCR

Total RNA was isolated and purified using the Quick-RNA Miniprep Kit (Zymo Research, Irvine, CA) according to the manufacturer’s protocol. RNA concentration was measured with The Qubit 2.0 Fluorometer (Thermo Fisher Scientific). RNA was reverse transcribed with the SuperScript IV First-Strand Synthesis System (Thermo Fisher Scientific, 18091050), using random hexamer primers provided within the kit. cDNA was amplified on an Applied Biosystems ViiA7 Real-Time PCR System with Power SYBR Green PCR Master Mix (Thermo Fisher Scientific, A25777). *Gapdh* was used as control. Data were analyzed using the ΔΔCt method unless otherwise stated. The following primer combinations were used: Ska2-fwd 5’ CCAAGAGCTGCATTTGTGCT 3’, Ska2-rev 5’ GGCTCTGTTGCAGCTTTCTC 3’; Gapdh-fwd 5’ TATGACTCCACTCACGGCAA 3’, Gapdh-rev 5’ ACATACTCAGCACCGGCCT 3’.

### Surgical procedures and viral injections

Mice were deeply anesthetized with ketamine/dexdormitor (medetomidine) mixture and placed in a stereotaxic apparatus (David Kopf Instruments, Tujunga, CA, USA). For virus delivery a 33-gauge microinjection needle with a 10-μl syringe (Hamilton) coupled to an automated microinjection pump (World Precision Instruments Inc.) at 2 nl/sec was used. Coordinates in millimeters from bregma were as follows [in mm]: A/P −1.9, M/L ± 1.3, D/V −1.8 and −1.3. At the end of the infusion, needles were kept at the site for 5 min and then slowly withdrawn. The injection volume was 0.5 µl. After bilateral infusion, incisions were sutured closed using nylon monofilament (Ethicon). During surgery, body temperature was maintained using a heating pad. After completion of surgery, anesthesia was reversed using Antisedan (atipamezole) and mice were allowed to recover on heating pads.

Surgeries for guide cannula implantations (microdialysis) were performed as previously described ^7,55^. Coordinates for microdialysis probe guide cannula implantations into the right mPFC were (with bregma as a reference point) as follows [in mm]: A/P 2.00, M/L 0.35, and D/V −1.50. After guide cannula implantation, animals were allowed to recover for 7 days in individual microdialysis cages (16 x 16 x 32 cm). Metacam (0.5 mg/kg, s.c) was injected within the first three days after surgeries, when required.

### Microdialysis

The perfusion setup consisted of lines comprised of FET tubing of 0.15 mm ID (Microbiotech Se, Sweden), a 15 cm-PVC inset tubing (0.19 mm ID), a dual-channel liquid swivel (Microbiotech Se, Sweden). Perfusion medium was sterile RNase free Ringer’s solution (BooScientific, USA) containing 1% BSA (Sigma, A9418). Perfusion medium was delivered to the probe at the flow rate of 0.38 μl/ min with the syringe pump (Harvard Apparatus, USA) and withdrawn with the peristaltic pump MP2 (Elemental Scientific, USA) at the flow rate of 0.4 μl/ min. Microdialysis CMA 12 HighCO Metal Free Probe was of 2 mm length membrane with 100 kDa cut off (Cat.N. 8011222, CMA Microdialysis, Sweden). All lines were treated with 5% polyethylenimine (PEI) for 16 h and then with H_2_O for 24 h before the experiments. The microdialysis probe was inserted into the implanted guide cannula (under 1-1.5 min isoflurane anesthesia, 2% in air) 6 days after the stereotaxic surgery and 18 h before the samples collection. A baseline sample collection phase (three samples) was always preceding the foot shock (FS), which allowed us to express the changes in the extracellular content of proteins as relative to the baseline values. On the experimental day, microdialysis fractions were constantly collected (for 30 min) on ice into 1.5 ml Protein LoBind tubes (Eppendorf) preloaded with 0.5 μl protease inhibitor cocktail 1:50 (Roche) at a perfusion rate of 0.4 μl/min. After collection, samples were immediately frozen on dry ice for subsequent expression analysis. After collection of three baseline samples animals were transferred to the FS chamber (ENV-407, ENV-307A; MED Associates, 7 St Albans, VT, USA) connected to a constant electric flow generator (ENV-414; MED Associates) and a FS (1.5 mA x 1 sec x 2) was delivered. After this procedure, mice were returned to the microdialysis cage where two post-FS samples were collected. To examine an effect of ULK1 inhibitor MRT 68921 on stress-evoked changes in extracellular content of proteins, the drug was injected intraperitoneally at a dose of 5.0 mg/kg and in a volume 10 ml/kg 4 h before the FS (the drug was prepared freshly dissolving a stock solution [60% EtOH/40% DMSO mixture] with saline at 1:20). At the end of the experiment, probes were removed, brains were frozen and kept at −80°C for the probe placement verification. 40 μm brain sections were stained with cresyl violet (Carl Roth GmbH) and probe placement was verified under a microscope. If probe placement was found to be out of the targeted region of interest, the respective samples were excluded from the study.

### Behavior

All experiments were analyzed using the automated video-tracking system ANYmaze (Stoelting, Wood Dale, IL).

## Open field (OF) test

The OF test was used to characterize locomotor activity in a novel environment. Testing was performed in an open field arena (50 x 50 x 50 cm) dimly illuminated (10 lux) in order to minimize anxiety effects on locomotion. All mice were placed into a corner of the apparatus at the beginning of the trial. The testing duration was 10 min and the distance traveled was assessed.

## Y-maze

The Y-maze test was used to assess spatial recognition memory. The apparatus consisted of three evenly illuminated arms (30 x 10 x 5 cm, 10 lux) with an angle of 120° between each arm. The apparatus was surrounded by various spatial cues. To ensure that the mice had sufficient spatial cues available without having to stretch up and look outside of the maze, we also introduced intra-maze cues (triangles, bars, and plus signs) that served the same purpose as the external cues. The Y-maze test comprises two trials separated by an intertrial interval (ITI) to assess spatial recognition memory. During the first trial, the mouse was allowed to explore only two of the three arms for 10 min while the third arm was blocked. After a 30 min ITI, the second trial was conducted during which all three arms were accessible for 5 min and the time spent in each arm was assessed. An increased amount of time spent in the novel arm, relative to the familiar arms, reflects intact spatial recognition memory.

## Novel object recognition memory task

Novel object memory was assessed in the Y-maze under low illumination (10 lux). During the acquisition trial, mice were presented with two identical objects and allowed to freely explore the objects for 10 min. Following a 30 min ITI, mice were presented with one familiar object and a novel one. During the retrieval phase mice were allowed to explore the objects for 5 min. At the start of each trial, mice were placed in the arm without an object. All objects were built from 10 LEGO bricks to allow a consistent volume, while shape and color could be varied to create distinguishable objects. The type of object that was chosen as familiar or novel respectively as well as the relative positions of the novel and familiar objects were counterbalanced across the groups. The time spent interacting with the objects was assessed and the ratio of time exploring the novel to the familiar object was calculated. A higher preference for the novel object reflects intact object recognition memory.

## Elevated plus maze (EPM)

The EPM was employed to assess anxiety-related behavior under low illumination (10 lux). The apparatus consisted of a plus-shaped platform with four intersecting arms, elevated 70 cm above the floor. Two opposing open (30 × 5 cm) and closed (30 × 5 × 15 cm) arms were connected by a central zone (5 × 5 cm). Animals were placed in the center of the apparatus facing the closed arm and were allowed to freely explore the maze for 5 min. Open arm time was calculated as a percentage of time in seconds: open arm time [%] = open arm time/(open arm time + closed arm time).

### Immunohistochemistry (mouse brain tissue)

Mice were deeply anesthetized with isoflurane and perfused intracardially with PBS followed by 4% paraformaldehyde. Brains were removed, post-fixed overnight in 4% paraformaldehyde followed by an additional overnight incubation in 30% sucrose solution at 4°C, and then stored at −80°C. Frozen brains were coronally sectioned in a cryostat microtome at 35 μm. Slices were subsequently washed with PBS and blocked using 10% normal donkey serum (NDS) prepared in PBS containing 0.1% Triton X-100 for 1 h at room temperature. Next, slices were incubated with the appropriate primary antibody (anti-NeuN, 1:1000, Synaptic Systems, 266004; anti-IBA1, 1:1000, FUJIFILM Cellular Dynamics, 019-19471; anti-GFAP, 1:1000, Cell Signaling Technology, 12389; anti-ASC, 1:200, AdipoGen, AG-25B-0006-C100; anti-CASP-1, 1:1000, Santa Cruz, sc-56036; anti-mCherry, 1:1000, Millipore Sigma, AB356481) in 10% NGS PBS overnight at 4°C on a shaker. Then slices were washed three times (10 min each) with PBS and incubated with the Alexa Fluor 488 and 594 conjugated secondary antibodies in 10% NGS PBS for 2 h at room temperature. Following three washes (15 min each) with PBS, slices were mounted on superfrost plus slides and covered with Vectashield mounting medium (Vector Laboratories, Burlingame, USA) containing DAPI. Slides were stored at 4°C until imaging.

### Imaging and quantification

Sixteen-bit images were acquired on a Leica SP8 confocal microscope with 10x or 40x objectives at identical settings for all conditions. Images were quantified using ImageJ (https://imagej.nih.gov/ij). For each experimental condition, one to two coronal sections per mouse from the indicated number of animals were used.

#### NeuN CA1 thickness

NeuN staining was used to measure the CA1 thickness with ImageJ. Leica SP8 with a 10x objective was used to acquire the images. The identical portion of the dorsal hippocampus was imaged for each brain.

#### Microglia

IBA1 immunoreactive cells were considered microglia. Leica SP8 with a 10x objective was used to acquire the images. The identical portion of the dorsal hippocampus was imaged for each brain. The cell counter plugin in ImageJ was used to count cells manually. When microglia density was too high to count individual cells, the signal intensity was measured in ImageJ instead.

#### Astrocytes

GFAP immunoreactive cells were considered astrocytes. Leica SP8 with a 10x objective was used to acquire the images. The cell counter plugin in ImageJ was used to count cells manually. When astrocyte density was too high to count individual cells, the signal intensity was measured in ImageJ instead.

#### CASPASE-1

Leica SP8 with a 10x objective was used to acquire the images. CASP-1 signal intensity was measured in ImageJ.

#### ASC

Leica SP8 with a 40x objective was used to acquire the images. The cell counter plugin in ImageJ was used to count ASC+ cells as well as ASC specks manually.

### Analysis of ASC specks in SIM-A9 cells

SIM-A9 wild type and SIM-A9 *Sec22b^-/-^* cells stably expressing ASC-mCerulean were used as reporter cells for inflammasome activation and generated as described before ^12^. For imaging experiments, SIM-A9 wild type and SIM-A9 *Sec22b^-/-^* cells expressing ASC-mCerulean were plated at a density of 2 × 10^5^ cells/well on black 96-well plates (µ-Plate, ibidi, Gräfelfing, Germany). Transfection of 200 ng of SCR- and Ska2-shRNA constructs was performed using Lipofectamine 3000 (Thermo Fisher Scientific) according to the manufacturer’s instructions. 48 h post transfection, cells were stimulated with 200 ng/ml LPS from *E. coli* 026:B6 (Thermo Fisher Scientific) for 2 h and subsequently fixed using 4% PFA. Images for assessment of ASC specks in PFA-fixed SIM-A9 cells were acquired using the VisiScope CSU-W1 spinning disk confocal microscope and the VisiView Software (Visitron Systems GmbH). Settings for laser and detector were maintained constant for the acquisition of each image. For analysis, at least seven images were acquired using the 20x objective. For quantification of ASC specks, mCerulean signal resembling an ASC speck/cell was counted manually in ImageJ and normalized to the number of DAPI- or DRAQ5-positive nuclei (ratio to cell count).

### Production of adeno-associated viruses (AAVs)

Packaging and purification of pAAV9-U6-shRNA[Ska2#1]-PGK-mCherry and pAAV9-U6-shRNA[Scr]-PGK-mCherry was conducted by Vigene Biosciences (Rockville, MD, USA). AAV9 titers were >1×10^13^ GC/ml. Packaging and purification of pAAV5-U6-shRNA[Ska2#2]-PGK-mCherry and pAAV5-U6-shRNA[Scr]-PGK-mCherry was conducted by the Viral Vector Core of Emory University (Altanta, GA, USA). AAV5 titers were >1×10^11^ GC/ml.

### Immunohistochemistry (human brain tissue)

Tissue blocks for immunohistochemistry were dissected from fresh brains and post-fixed in 0.1M phosphate buffer (PB) containing 4% paraformaldehyde and 0.1M NaN_3_ at 4°C for 3 weeks, then cryoprotected at 4°C (30% glycerol, 30% ethylene glycol and 0.1% NaN_3_ in 0.1M PB), embedded in agar, and pre-sliced in 2.5 mm coronal slabs using an Antithetic Tissue Slicer (Stereological Research Lab., Aarhus, Denmark). Each slab was exhaustively sectioned using a freezing microtome (American Optical 860, Buffalo, NY). Sections were stored in cryoprotectant at −20°C. Sections were cut at 50 µm thickness through the hippocampus and collected in compartments in serial sequence. Four to six sections within one compartment/subject were selected for immunolabeling.

## Immunocytochemistry

Antigen retrieval was carried out by placing free-floating sections in citric acid buffer (0.1 M citric acid, 0.2 M Na_2_HPO_4_) heated to 80°C for 30 min. Sections were then incubated in primary antibody (SKA2, SAB3500102, lot#54031701, MilliporeSigma, St. Louis, MO) for 48 h at 4°C and then in biotinylated secondary serum (SKA2, goat anti-rabbit IgG 1:500; Vector Labs, Inc. Burlingame, CA). This step was followed by streptavidin conjugated with horse-radish peroxidase for two h (1:5000, Zymed, San Francisco, CA), and, finally, nickel-enhanced diaminobenzidine/peroxidase reaction (0.02% diaminobenzidine, Sigma-Aldrich, 0.08% nickel-sulphate, 0.006% hydrogen peroxide in PB). All solutions were made in PBS with 0.5% Triton X unless otherwise specified. All sections were mounted on gelatin-coated glass slides, coverslipped and coded for quantitative analysis blinded to age. Sections from all brains included in the study were processed simultaneously within the same session to avoid procedural differences. Each six-well staining dish contained sections from normal control subjects and was carried through each step for the same duration of time, so to avoid sequence effects. Omission of the first or secondary antibodies did not result in detectable signal.

## Dual antigen immunofluorescence

Antigen retrieval as described above. Sections were co-incubated in primary antibodies (rabbit anti-SKA2, 1:300, SAB3500102, MilliporeSigma, St. Louis, MO; mouse anti-IBA1, 1:500, cat# 013-27593, Wako FujiFilm Chemicals USA Corp., Richmond, VA; mouse anti-CamKIIα 1:500, ab22609, Abcam, Cambridge, MA) in 2% BSA for 72 h at 4°C. This step was followed by 4 h incubation at room temperature in Alexa Fluor goat anti-mouse 647 (1:300; A-21235, Invitrogen, Grand Island, NY) and donkey anti-rabbit 488 (1:300; A-21206, Invitrogen, Grand Island, NY), followed by 1 min incubation in TrueBlack solution (cat# 23007, Biotum Inc., Fremont, CA) to block endogenous lipofucsin autofluorescence ^56^. Sections were mounted and coverslipped using Dako mounting media (S3023, Dako, North America, Carpinteria, CA).

## Data collection

An Olympus BX61 interfaced with StereoInvestigator v.2019 (MBF Biosciences, Willinston, VT) was used for analysis. The borders of the hippocampal subregions were identified according to cytoarchitectonic criteria as described in our previously published studies ^57,58^. A 1.6x objective was used to trace the borders of hippocampal subregions. Each traced region was systematically scanned through the full x, y, and z axes using a 40x objective to count each immunoreactive (IR) cell within the traced borders over all sections from each subject.

## Numerical densities and total numbers estimates

Numerical densities (Nd) were calculated as *Nd =* Σ*N /* Σ*V* where *N* is the sum of cells within a region of interest, and V is the total volume of the region of interest as described previously in detail ^59^.

### PheWAS

Phenotypic data for the *FKBP5* and *SKA2* genes were obtained from the Atlas of GWAS Summary Statistics (http://atlas.ctglab.nl/PheWAS), database release3: (v20191115) ^16^

### Statistical analysis

The data presented are shown as means + standard error of the mean (SEM). Cell culture and mouse data were analyzed using GraphPad 7.0 (La Jolla, CA). When two groups were compared, paired or unpaired, two-tailed Student’s t-test was applied, as appropriate. For three or more group comparisons, one-way, two-way or repeated measures two-way analysis of variance (ANOVA) was performed, followed by Tukey’s, Bonferroni or Šidák’s multiple comparison post hoc test, as appropriate. JMP Pro v. 14 SW (SAS Institute Inc., Cary, NC) was used for analysis of covariance (ANCOVA) of human postmortem data. Differences between groups relative to the main outcome measures in each of the regions examined were assessed for statistical significance using an ANCOVA stepwise linear regression process. Age, sex, and postmortem time interval, hemisphere, and brain weight are included in the model if they significantly improved the model goodness-of-fit.

P-values of < 0.05 were considered statistically significant.

### Data and code availability

Original/source data for Figure S2A and B (Phenotypic data for the *FKBP5* and *SKA2)* genes were publicly available from the Atlas of GWAS Summary Statistics and can be downloaded at http://atlas.ctglab.nl/PheWAS.

## Acknowledgments

We thank all lab members for suggestions and comments on the experiments and manuscript. This study was supported by a NARSAD Young Investigator Grant from the Brain & Behavior Research Foundation, honored by P&S Fund (awarded to N.C.G., Grant ID 25348), the National Institutes of Health (awarded to K.J.R., P50-MH115874 and R01-MH117292). T.K. was supported by research grants from NICHD (R21HD088931, R21HD097524, R01HD102974), NIMH (R21MH117609), and ERA-Net Neuron (01EW2003). Human tissue was obtained from the NIH NeuroBioBank. In addition, tissue samples were supplied by The Manchester Brain Bank, which is part of the Brains for Dementia Research programme, jointly funded by Alzheimer’s Research UK and Alzheimer’s Society. Fig. 1O, 1P and 3L were created with BioRender.

## Author contributions

J.H., K.J.R. and N.C.G. conceived the project and designed the experiments. T.B., C.K. K.H., T.R., S.M. and N.C.G. performed cell culture experiments. J.H., C.K., A.K.G., E.A., F.T., G.M., M.L.P., D.E.H., J.O., R.L., T.K., N.D, K.M.M., T.P., V.S., E.L., W.A.C. and M.V.S. performed animal experiments. J.H., C.K., L.R., K.L., A.P., A.C.R., C.M., S.B., T.K., H.P. and N.C.G. performed human postmortem experiments. J.H., K.J.R. and N.C.G wrote the initial version of the manuscript. J.H., K.J.R. and N.C.G supervised the research. All authors contributed to the final version of the manuscript.

## Supplementary Information

### Supplemental Figures and Legends

**Figure S1.**
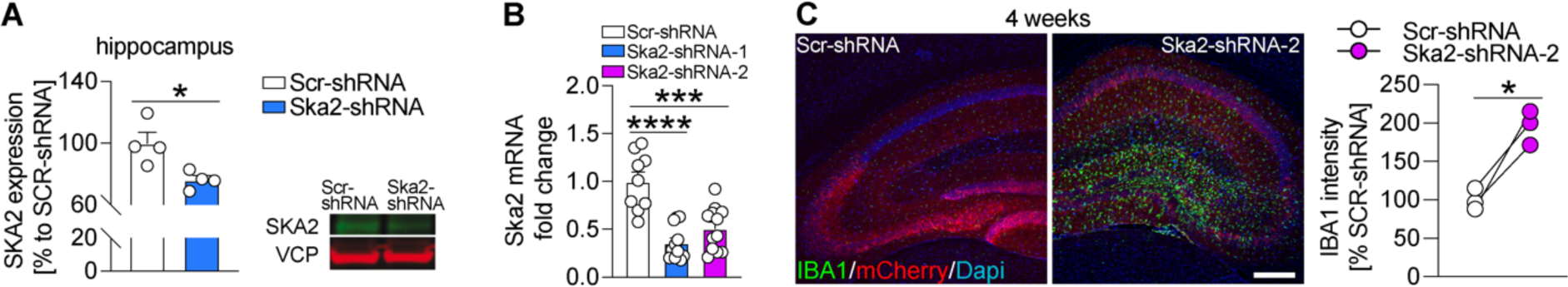
**(A)** Viral-mediated knockdown of *Ska2* (Ska2-shRNA-1-AAV) in the hippocampus leads to significantly decreased SKA2 expression (n = 4 per group). **(B)** Ska2 mRNA expression is significantly decreased following transcfection with Ska2-shRNA-1 or Ska2-shRNA-2 in Neuro2a cells. **(C)** Viral-mediated knockdown of *Ska2* (Ska2-shRNA-2-AAV) leads to increased IBA1 expression 4 weeks after viral injection. Unpaired, two tailed t-test for simple comparisons, one-way analysis of variance (ANOVA) + Tukey’s post hoc test or paired t-test: * = p < 0.05; *** = p < 0.001; **** = p < 0.0001. Scale bar represents 250 µm.

**Figure S2.**
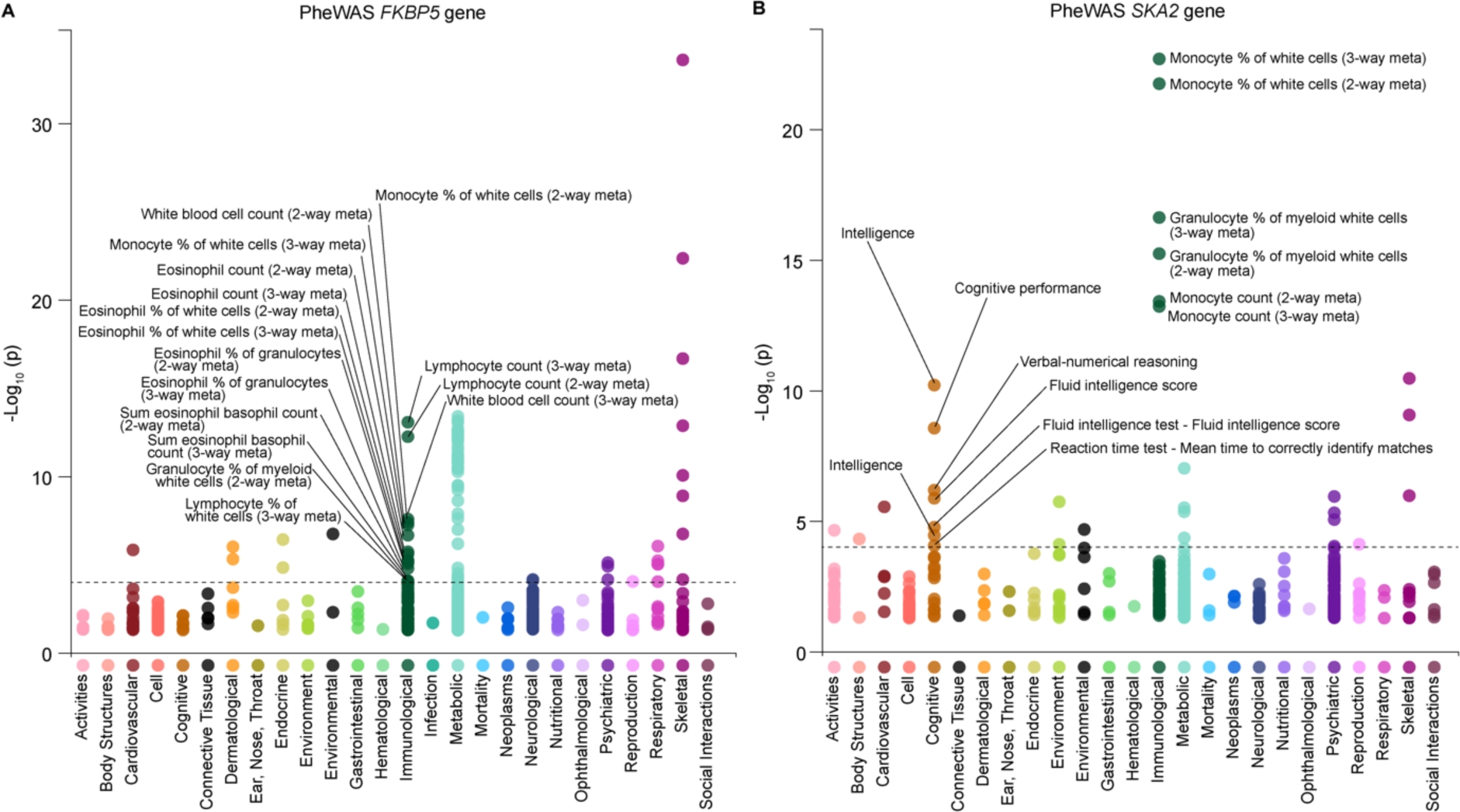
PheWAS plots of phenotypes associated with the *FKBP5* (A) and *SKA2* (B) gene. The x axes represent phenotypes, and the y axes represent the −log_10_ of uncorrected p values. The dashed lines indicate the experiment-wide threshold to survive Bonferroni correction (*FKBP5*: p_-Log10_ < 4.016 and *SKA2*: p_-Log10_ < 3.876). Each dot represents one phenotype, and the colors indicate their according traits. Representative top findings are annotated in the figure.

**Table S1.** Phenome-Wide Association Studies (PheWAS) table of the *FKBP5* locus.

**Table S2.** Phenome-Wide Association Studies (PheWAS) table of the *SKA2* locus.

**Table S3.** Details of human postmortem subjects (Immunoprecipitation).

**Table S4.** Details of human postmortem subjects (Immunohistochemistry).

**Table S5.** Details of human postmortem subjects (Alzheimer’s disease discovery cohort (HIP)).

**Table S6.** Details of human postmortem subjects (Alzheimer’s disease replication cohort (PFC)).

